# VCP is a critical component for TDP-43 disaggregase activity in skeletal muscle

**DOI:** 10.64898/2026.09.14.751559

**Authors:** Eileen M Lynch, Sara Pittman, Jil Daw, Conrad C. Weihl

## Abstract

TDP-43 aggregates are a unifying pathological feature between several neurodegenerative diseases and myopathies. Mutations in the AAA ATPase protein VCP have been linked to TDP-43 proteinopathies in both brain and skeletal muscle including frontotemporal dementia (FTD), amyotrophic lateral sclerosis (ALS), and inclusion body myopathy (IBM). In the case of multisystem proteinopathy (MSP), patients with VCP mutations may present with combinations of these phenotypes. Of these MSP phenotypes IBM is the most common, affecting around 90% of patients. However, it is not well understood how mutations in VCP contribute to the accumulation of insoluble TDP-43 aggregates in the context of skeletal muscle. To study this further, we used *in vitro* and *in vivo* models of TDP-43 aggregation in skeletal muscle in the presence of VCP mutation or inhibition to study its effects on the ability to clear TDP-43 aggregates. Across multiple model systems we found that VCP disease mutations or inhibition causes a reduced capacity to clear insoluble TDP-43, suggesting a VCP loss of function in patients with VCP-related MSP.

## Introduction

Valosin containing protein (VCP, or p97) is a ubiquitously expressed hexameric protein of the AAA+ family (ATPases Associated with diverse cellular Activities) [1]. It is classified as a segregase as it uses the energy provided from ATP to extract substrate proteins from macromolecular complexes, then to be passed on for degradation through the ubiquitin proteasome or autophagy systems [2]. Mutations in VCP have been linked to a number of patient phenotypes including frontotemporal dementia (FTD), amyotrophic lateral sclerosis (ALS), inclusion body myopathy (IBM), and Paget’s disease of the bone (PDB) [3-6]. This spectrum of diseases caused by mutations in VCP and others such as hnRNPA2B1, hnRNPA1, SQSTM1, MATR3 is now referred to as multisystem proteinopathy (MSP) [7]. Within VCP-associated MSP, 90% of patients present with myopathy, 42% with PDB, 30% with FTD, and 9% with ALS (with varying combinations of each) [8]. Notably, IBM is the most prevalent presentation.

Aggregates of the RNA-binding protein TDP-43 are a common pathological feature in both neurons and skeletal muscle of patients with MSP. In healthy cells TDP-43 is primarily localized to the nucleus where it has several roles in RNA processing. However, it does shuttle between the nucleus and cytoplasm under certain conditions, joining temporary cytoplasmic membrane-less structures such as stress granules, RNA transport granules, and myo-granules [9, 10]. Through mechanisms that have yet to be completely characterized, in some cases TDP-43 remains in the cytoplasm and forms more permanent insoluble aggregates. TDP-43 aggregates can also form inside nuclei, most notably in neurons [11]. Aside from toxicity caused by the aggregates themselves, this sequestration of TDP-43 also causes a loss of function effect through disrupted RNA processing. In particular, one notable role of TDP-43 is to regulate the splicing of cryptic exons. The abnormal inclusion of these normally excluded exons results in either nonsense mediated decay of the transcript or a novel protein product [12, 13].

TDP-43 aggregation and dysfunction appears to be a critical driver of pathology in both neurodegeneration and myopathy. However, how mutations in VCP lead to TDP-43 aggregation is not fully understood, particularly in skeletal muscle. With such broad cellular functions, VCP could feasibly influence TDP-43 pathology at the level of aggregate accumulation, clearance, or spread, through many different mechanisms such as alterations to the ubiquitin-proteasome system, autophagy, vesicle trafficking, etc. In particular, VCP is known to be involved in stress granule clearance [14-16]. While TDP-43 is a component of stress granules, the relationship between temporary stress granule-like structures and more permanent TDP-43 aggregates is not fully understood. There are likely many intermediate stages of TDP-43 oligomerization, liquid-liquid phase separation, and aggregation which may contribute to cellular pathology in different ways. Therefore, we employed a variety of *in vitro* and *in vivo* systems to study the relationship between VCP and the clearance of insoluble TDP-43. *In vitro*, we found that VCP inhibition results in impaired clearance of insoluble TDP-43 from a cell line with inducible expression of TDP-43^ΔNLS^, arsenite treated myocytes, and TDP-43 FRET biosensor cells that have been seeded with TDP-43 aggregates. *In vivo*, we crossed HSA-TDP-43^ΔNLS^ mice with VCP^R155H/WT^ knock in mice and again found impaired clearance of TDP-43 aggregates in the VCP mutant mice. Together these experiments support a loss of function role of VCP in TDP-43 proteinopathy through a reduced capacity for returning TDP-43 to its soluble state.

## Materials & Methods

### C2C12 culture

Immortalized mouse myoblast C2C12 cells (American Type Culture Collection, catalog no. CRL-1772, RRID:CVCL_0188) were maintained in Dulbecco’s modified Eagle’s medium (DMEM) with 10% FBS and 0.05% penicillin and streptomycin (Thermo Fisher, 15140122) and passaged using 0.05% trypsin-EDTA. Once 90 percent confluent, the cells were switched to a low serum medium of DMEM and 2% horse serum, with daily media changes for 5-8 days before treatment.

### Primary myoblast culture

Both human and mouse primary myoblast lines were cultured on Matrigel coated plates. A 1:50 solution of Matrigel (Corning, 356237) was added to the plates and incubated at room temperature for 1 hour then removed before use. Cells were cultured in proliferation medium containing 20% fetal bovine serum (FBS; GeminiBio, 100-800), 1% penicillin and streptomycin, 0.25 µg/mL amphotericin B (Gibco 15290-018), 1% MEM non-essential amino acids (Gibco 11140), and 0.001% β-mercaptoethanol (Sigma-Aldrich, M6250) in DMEM/F12 (Gibco, 11320033). Basic fibroblast growth factor (bFGF; R&D Systems, 233-FB) was added to the media at a final concentration of 10ng/mL before use. Once cells were confluent, differentiation was induced by switching to a lower serum differentiation medium consisting of 5% horse serum (Gibco, 16050122) and 1% penicillin/streptomycin in DMEM with daily media changes.

### Arsenite treatment

Cells were treated with 1 mM arsenite for 1 hour then washed with PBS and replaced with regular media for recovery. At specific recovery timepoints, cells were washed with PBS, aspirated, and flash frozen in -80 until ready to be lysed in RIPA buffer and processed for BCA.

### Soluble/insoluble fractionation

Flash frozen cell culture plates were scraped and mouse muscle samples were homogenized in RIPA buffer with protease inhibitors (Sigma-Aldrich, S8820). Samples were sonicated with 10 cycles of 30 s on, 30 s off at amplitude 50, then centrifuged at 13,000 rpm for 10 minutes at 10 °C to pellet the debris. BCA assay was used to get supernatant protein concentrations and samples were diluted to equal starting concentrations. The samples were again sonicated before beginning the fractionation process. The RIPA insoluble fraction was pelleted by ultracentrifugation at 100,000 xg and 4 °C for 30 minutes. The supernatant was saved as the soluble fraction and the pellet was rinsed with RIPA buffer, sonicated, and spun down again at the same speed and conditions. The final pellet was resuspended in 7 M urea buffer with 2 M thiourea, 4% CHAPS, and 30 mM tris (pH 8.5). If preparing samples for use in the FRET assay, lysates are rinsed with PBS instead of RIPA, and the final pellet is resuspended in PBS as well.

### FRET biosensor line culture and seeding

The TDP-43 FRET sensor line consists of HEK293 cells stably expressing c-terminal fragments (aa262-414) of TDP-43 tagged with mRuby3 and mClover3 at the c-terminal end. The cell line was maintained in medium containing 10% fetal bovine serum and split using 0.05% trypsin. The cells were plated in 96-well plates at 35,000 cells per well to set up for seeding assays. After 24 hours, the cells are around 50% confluent and ready for treatment with TDP-43 seeds. The PBS-resuspended RIPA-insoluble fraction from cell or tissue lysates was incubated with Lipofectamine 2000 transfection reagent (1 µL per well, Thermo Fisher Scientific, 11668019). After 72 hours of incubation with seeds, the cells were fixed to be analyzed by flow cytometry. To fix, cells were trypsinized, transferred to a round-bottom 96-well plate and centrifuged at 1,000 xg for 5 minutes. Then cells were resuspended in 2% paraformaldehyde for 10 minutes, centrifuged again, and resuspended in 150 µL of flow running buffer and stored at 4°C until ready for use.

Cells were analyzed for aggregation using a MACSQuant VYB flow cytometer. A 488-nm laser and 525/50-nm band-pass filter was used to detect mClover3 signal, the 561-nm laser and 615/20-nm filter were used to detect mRuby3 signal, and FRET was measured through 488-nm laser excitation and 614/50-nm filter. Integrated FRET density was calculated as the percent positive cells multiplied by the median fluorescence intensity of the FRET channel, normalized to a buffer only control.

### Western blotting

Cells or tissue were lysed in RIPA buffer with protease and phosphatase inhibitors. Protein concentrations were determined using a BCA assay and measured with the BioTek Synergy H1 plate reader (Agilent) at a wavelength of 562 nm. Samples were diluted to equal concentrations, mixed with loading buffer and boiled for 5 minutes at 95 °C before loading into 10% SDS-PAGE gels at 20-40 µg of total protein per lane, or equal volumes of soluble/insoluble samples. Gels were run at 100 V for 10 minutes then 120 V for 1 hour and 15 minutes. The samples were transferred to nitrocellulose membranes for 2 hours at 26 mA. Membranes were blocked with 5% milk in PBS-T for 1 hour. Primary antibodies were diluted according to **Table 1** in 5% milk PBS-T and incubated with the membranes overnight at 4 °C. The primary antibodies were washed off by three PBS-T washes of 5 minutes each. The membranes were then incubated with anti-rabbit or anti-mouse horseradish peroxidase (HRP) secondary antibodies in 5% milk PBS-T at a 1:5000 dilution for 1 hour at room temperature, then washed off with three PBS-T washes. Membranes were imaged using enhanced chemiluminescence (ECL; Bio-Rad, 1705060) and the Syngene G:Box Chemi XT4 imager.

**Table 1.** Information on the primary antibodies used in western blotting.

| Marker | Catalog # | Dilution |
| --- | --- | --- |
| TDP-43 | Proteintech 10782-2-AP | 1:1000 |
| pTDP-43 | Proteintech 22309-1-AP | 1:1000 |
| GAPDH | Cell Signaling 2118 | 1:1000 |
| VCP | BD 612182 | 1:500 |
| P62 | Abnova H0008878 | 1:1000 |

### Immunocytochemistry

Cells were plated onto Matrigel coated coverslips. Cells were fixed with ice cold methanol for 10 minutes followed by three 5 minute washes with 1x PBS. Blocking/permeabilization was performed with 5% NGS and 0.1% Triton X-100 in PBS. Primary antibodies were diluted in 5% NGS and 0.1% Triton X-100 in PBS according to the dilution factors in **Table 2** and incubated with the cells for one hour at room temperature followed by another three 1x PBS washes. Secondary antibodies were diluted 1:1000 in 5% NGS in PBS and incubated for 30 minutes at room temperature followed by another three 1x PBS washes. Cells were stained for 10 minutes with 4’,6-diamidino-2-phenylindole (DAPI; 1 µg/mL) followed by the final three 1x PBS washes. Coverslips were mounted onto slides using Mowiol 4-88 (Sigma-Aldrich, 81381).

**Table 2.**
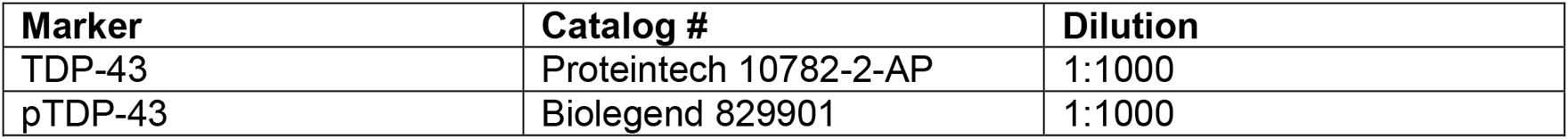
Information on the primary antibodies used in immunocytochemistry.

### Histochemistry/immunohistochemistry

Following dissection, mouse hind limb muscles were mounted onto corks using 10% tragacanth gum (Sigma-Aldrich, G1128) and flash frozen in 2-methylbutane over liquid nitrogen before storing at -80 °C. The frozen muscle was sectioned at 10 µm thickness. H&E stains were performed with a 1% aqueous solution of eosin Y (Sigma-Aldrich, E-6003) and Harris hematoxylin stain (Lerner Laboratories, 1931382), filtered before use. Slides were immersed in the Harris hematoxylin for 10 s then transferred to a beaker of tap water and rinsed until the water was clear. Then slides were immersed in eosin stain for 30 s and rinsed with tap water as before. The sections were then dehydrated via ascending alcohol solutions (50%, 70%, 80%, 95% x 2, and 100% x2) and cleared with xylene three to four times. A glass coverslip was mounted over the sections using Permount mounting medium (Fisher, SP15-100).

For immunostaining, the mouse muscle sections were fixed for 10 minutes with 3.7% paraformaldehyde (PFA), washed three times with 1x PBS followed by 10 minutes of ice-cold acetone. Sections were permeabilized with 0.5% Triton-X 100 for 20 minutes then blocked with PerkinElmer blocking reagent (FP1012) for 1-2 hours at room temperature. Sections were incubated for either one hour at room temperature or overnight at 4 °C in primary antibodies in blocking reagent at dilutions listed in **Table 3**. The primary antibodies were washed off with three rinses of 1x PBS for 5 minutes each. Then, secondary antibodies were added at a 1:1000 dilution factor in blocking reagent and incubated for 1 hour at room temperature. Slides were again washed three times with 1x PBS. For nuclear stains, the slides were incubated for 10 minutes with 4’,6-diamidino-2-phenylindole (DAPI; 1 µg/mL) followed by another three 1x PBS rinses. Finallly, cover class was mounted onto the slides using Mowiol 4-88.

**Table 3.** Information on the primary antibodies used in immunohistochemistry.

| Marker | Catalog # | Dilution |
| --- | --- | --- |
| TDP-43 | Proteintech 10782-2-AP | 1:1000 |
| pTDP-43 | Proteintech 22309-1-AP | 1:1000 |
| Caveolin-3 | Thermo Fisher PA1-066 | 1:250 |

Slides were imaged using a Leica DM4 B upright microscope with THUNDER imaging technology and the Leica Application Suite X software.

### Mouse studies

As previously outlined [17], HSA-hTDP-43^ΔNLS^ mice were generated by breeding HSA-rtTA mice (Jackson Laboratory, strain 012433) with tetO-hTDP-43^ΔNLS^ mice (Jackson Laboratory, strain 014650). Breeding pairs were also established with VCP^R155H/WT^ knock in mice [18] until genotyping confirmed mice heterozygous for the knock in VCP mutation and both transgenes. All mice were bred on a C57BL/6 background and genotyped by Transnetyx. To activate the transgene, mice were treated with a doxycycline chow diet (200 mg/kg, Fisher Scientific, 14-727-450). Both male and female mice were used in experiments in equal numbers as much as possible, without any noted differences. Animal housing and all procedures were in accordance with protocols approved by the Animal Studies Committee at Washington University in St Louis.

### Statistical Analysis

GraphPad Prism was used to analyze all quantitative data. Graphs are presented as the mean ± SEM. All quantitative experiments contained at least three technical replicates. Biological replicates can be found in the figure legends. For multi-condition data sets, one-way analysis of variance (ANOVA) followed by Dunnett’s multiple comparisons test was used to detect significance. If only two groups were being compared, a two-tailed unpaired Student’s t test was used. For all experiments, \**P* < 0.05, \*\**P* < 0.01, and \*\*\**P* < 0.001.

## Results

### VCP inhibition slows the clearance of cytoplasmic mislocalized TDP-43 in a stable inducible cell line

First, we utilized a HEK293 cell line with tetracycline inducible expression of mcherry tagged hTDP-43^ΔNLS^ [19]. A time course was established in which expression was induced with tetracycline treatment for 48 hours and then cells were rinsed with PBS and returned to tetracycline-free medium. Samples were collected daily and lysed with RIPA buffer and the RIPA-insoluble fraction was isolated and analyzed by western blot to determine the clearance rate of insoluble TDP-43. We found that most of the insoluble TDP-43 was cleared from these cells by 2 days of recovery (**Fig 1A**). Using this time course, we next added VCP inhibitor CB-5083 during the recovery period following the washout of tetracycline. There was a persistence of insoluble TDP-43 at 24 and 48 hours following washout, whereas the insoluble TDP-43 was mostly cleared in the control cells which did not receive CB-5083 (**Fig. 1B**). This supports that VCP has an important role in the clearance of insoluble TDP-43.

**Figure 1.**
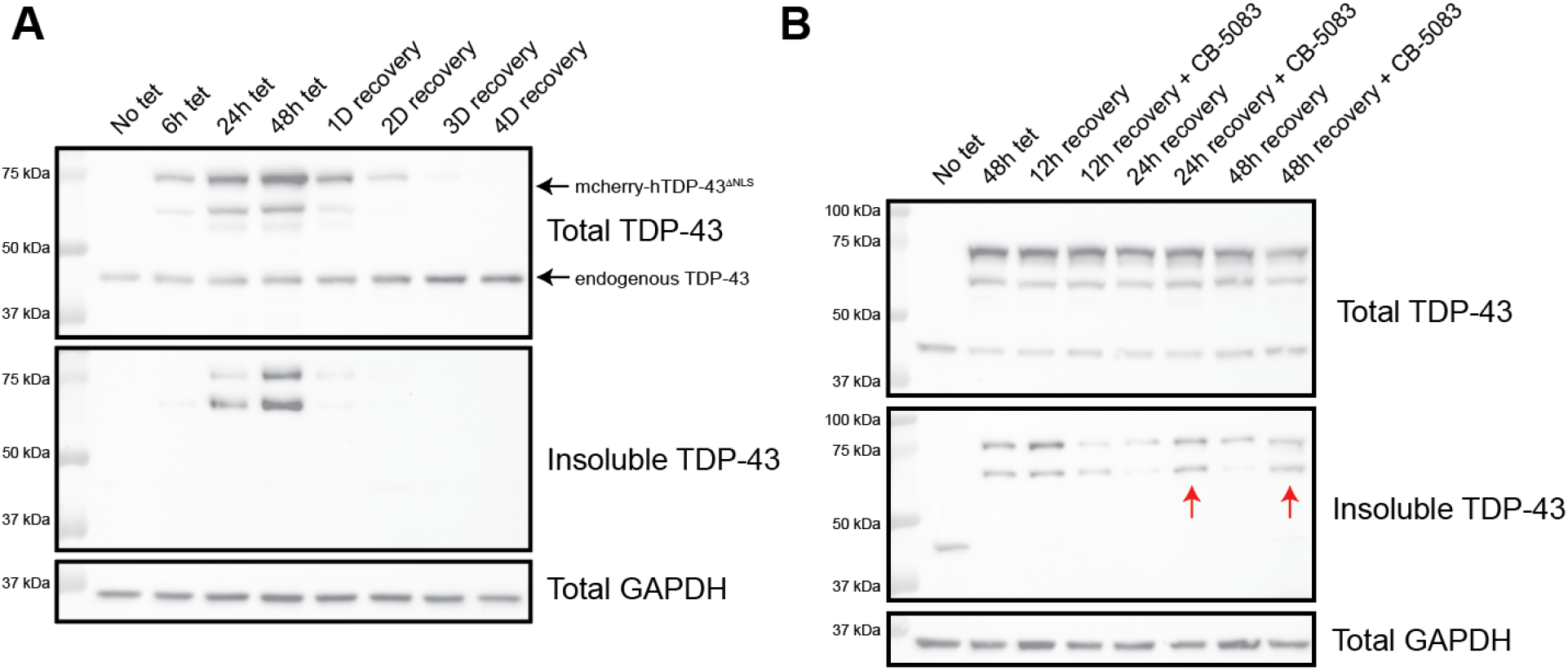
VCP inhibition delays clearance of insoluble TDP-43 in an inducible cell model. **(A)** HEK293 cells stably expressing tetracycline-inducible mcherry-TDP-43^ΔNLS^ were treated with 1 μg/mL of tetracycline for 48 hours then rinsed with PBS and returned to regular growth medium and collected at specific times to analyze levels of insoluble TDP-43 which was mostly cleared by 2 days of recovery. **(B)** A similar time course of 48 hours treatment with tetracycline followed by washout was performed and samples were collected at specific recovery time points with and without the incubation with 5 μM CB-5083. Cells treated with CB-5083 had a delayed clearance of mcherry-tagged TDP-43 (red arrows).

### VCP inhibitors slow the clearance of arsenite induced insoluble TDP-43 in C2C12 myotubes

To then test the effects of VCP inhibition on insoluble TDP-43 clearance specific to muscle cells, we utilized C2C12 cells, an immortalized mouse myoblast line. C2C12 myoblasts were differentiated into myotubes for 5 days and then treated with 1 mM sodium arsenite for 1 hour to induce the formation of insoluble TDP-43. The arsenite was then removed, the cells were rinsed with 1x PBS and returned to regular differentiation medium. Myotubes were lysed at specific recovery time points following arsenite treatment and analyzed for RIPA insoluble TDP-43. In this cell model, the insoluble TDP-43 was mostly cleared by 8 hours of recovery (**Fig. 2A**). Using this timeline, C2C12 cells were again treated with arsenite followed by 4 and 8 hours of recovery with or without VCP inhibitor CB-5083. Again we saw a persistence of insoluble TDP-43 in the CB-5083 compared to untreated controls (**Fig 2B**). Interestingly, VCP was in the insoluble fraction during the clearance time points but greatly reduced in the presence of CB-5083. We hypothesize that the VCP inhibitor may be preventing VCP from interacting with insoluble TDP-43. This model is valuable as it represents muscle-specific processing of endogenous insoluble TDP-43 rather than an overexpression of mutated TDP-43 as in the previous model.

**Figure 2.**
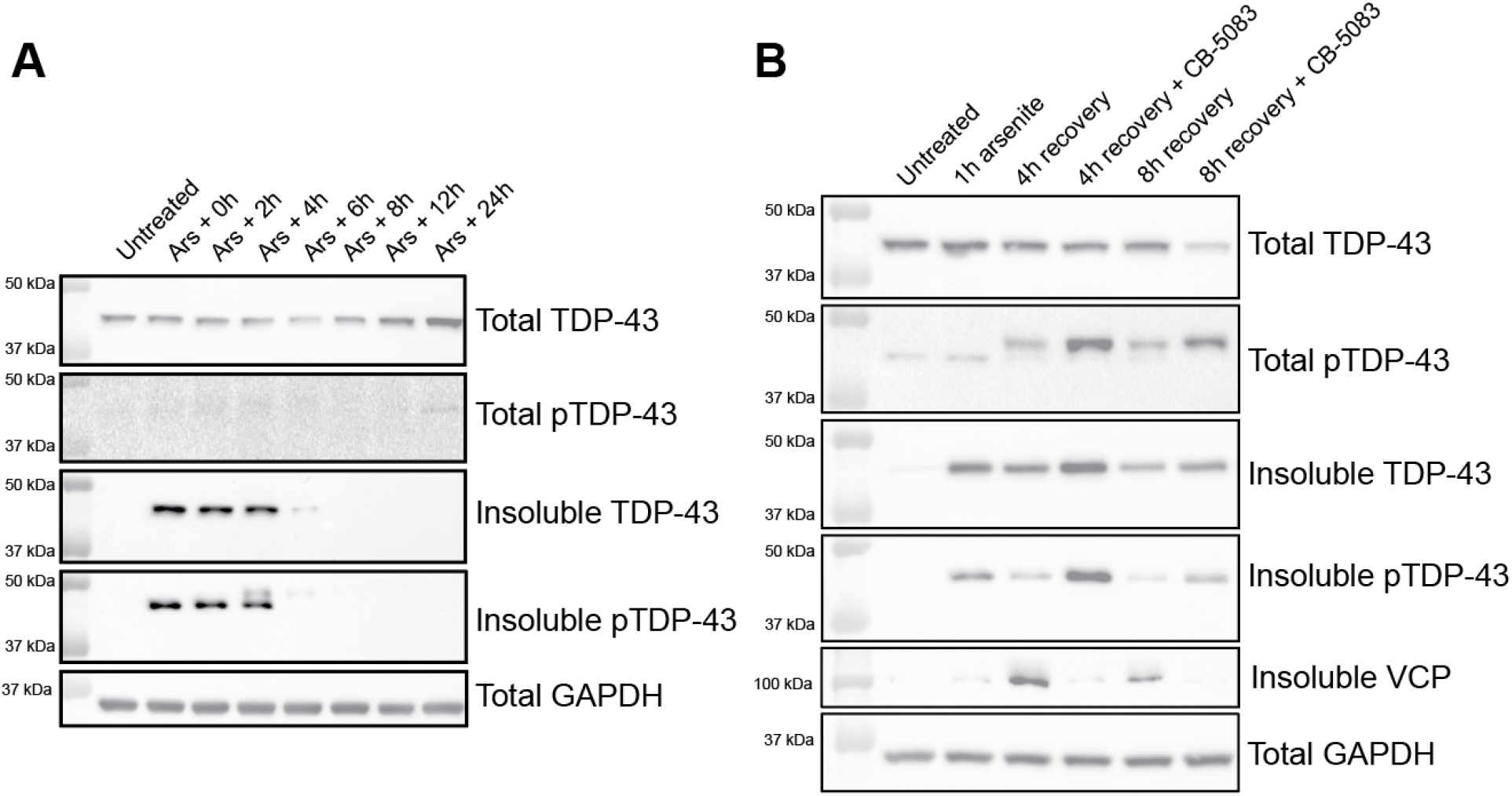
VCP inhibition slows the clearance of arsenite-induced insoluble TDP-43 in cultured myotubes. **(A)** C2C12 differentiated myotubes were treated with 1 mM sodium arsenite for 1 hour then washed with PBS before being replaced with regular medium and collected at specific recovery timepoints. Insoluble TDP-43 forms immediately following arsenite treatment and persists until around 8 hours. **(B)** The addition of 5 µM CB-5083 delays the clearance of insoluble TDP-43 following arsenite treatment.

**Figure 3.**
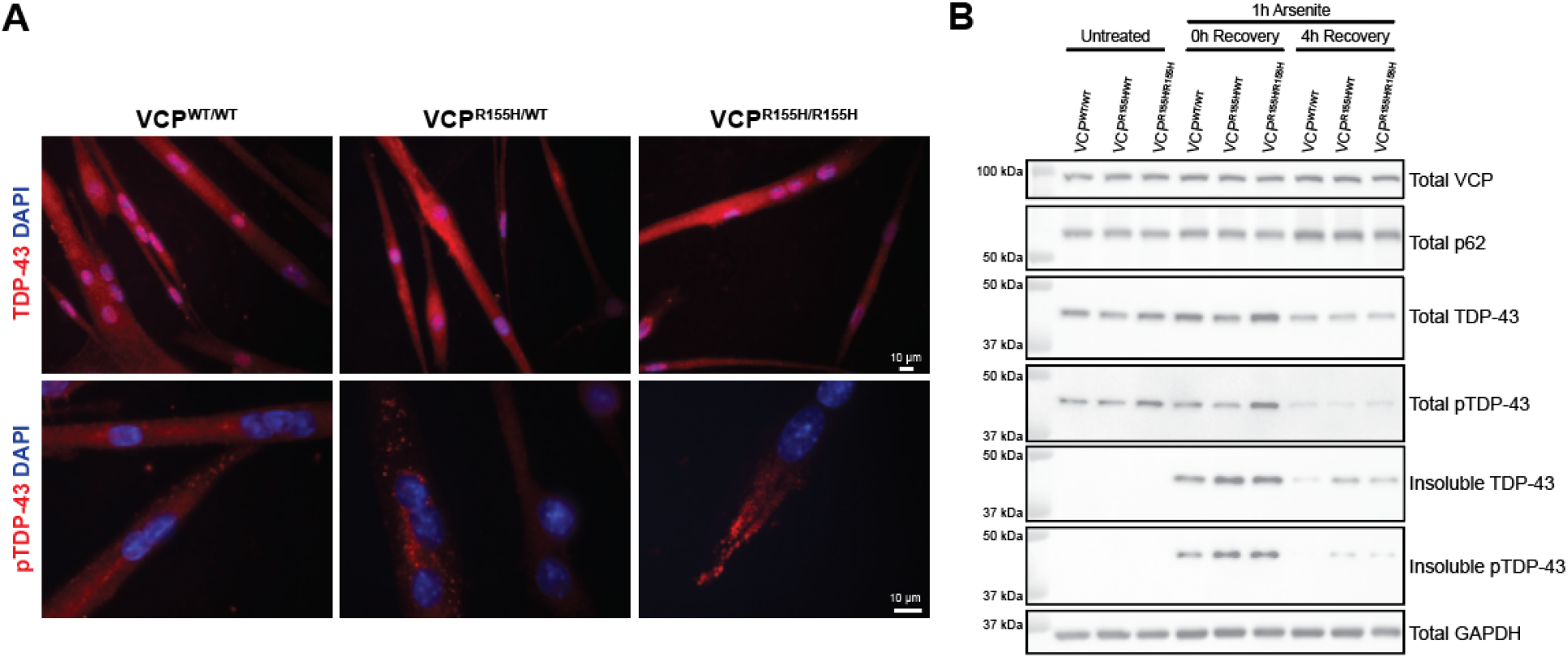
VCP mutations also delay clearance of insoluble TDP-43 in mouse primary myoblast lines. **(A)** Representative images of primary myoblast lines derived from WT, VCP-R155H heterozygous mice, and VCP-R155H homozygous knock in mice, with increased aggregate-like pTDP-43 in the mutant lines at baseline without any treatments. **(B)** While WT myocytes were able to largely clear insoluble TDP-43 by 4 hours post arsenite treatment, the heterozygous and homozygous VCP mutant lines had persistent insoluble TDP-43.

**Figure 4.**
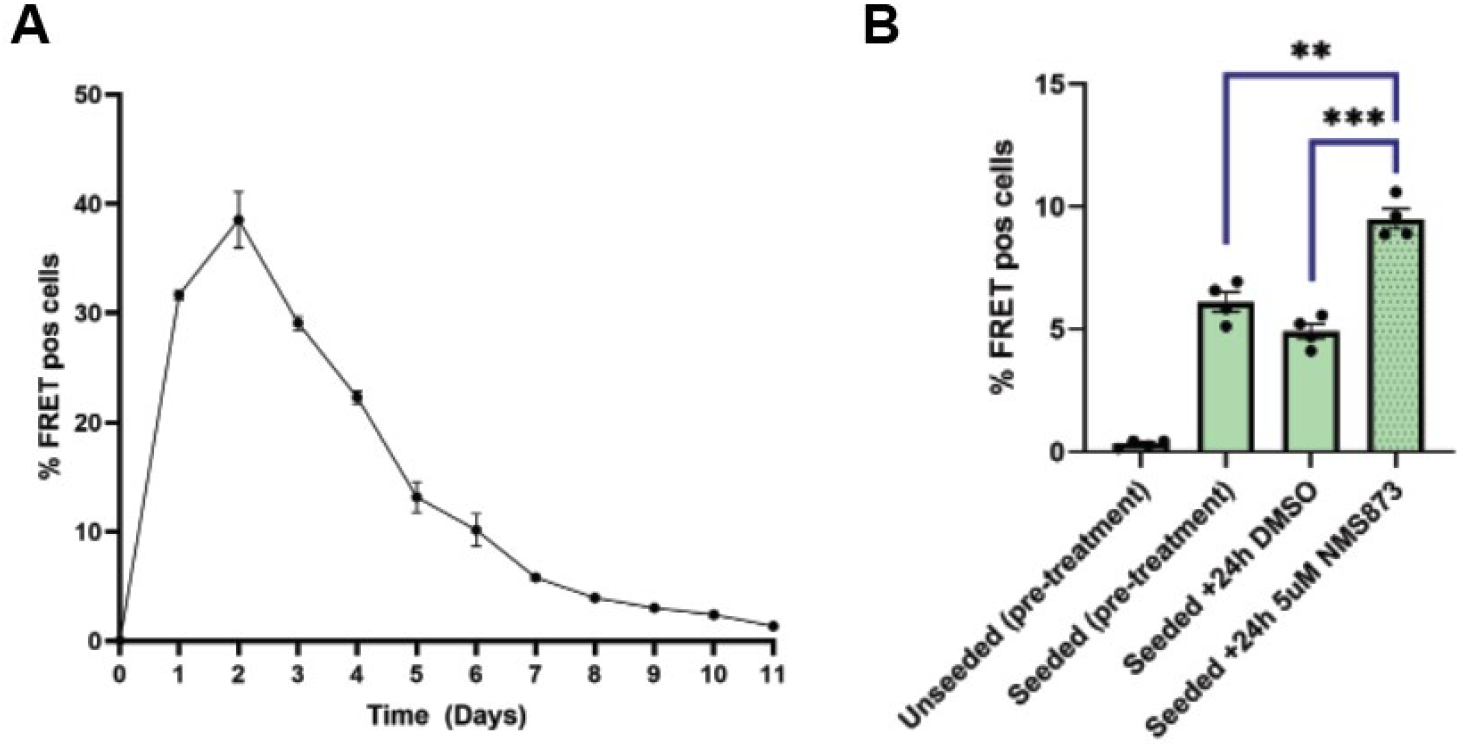
Seeded TDP-43 FRET biosensor cells have decreased clearance of TDP-43 aggregates in the presence of VCP inhibition. **(A)** Amount of FRET positive cells indicating aggregate formation and clearance over time. **(B)** Quantification of percentage of cells positive for FRET signal indicating TDP-43 aggregates, before and after a 24 hour treatment with 5 µM NMS-873.

### VCP mutations slow the clearance of arsenite-induced insoluble TDP-43 in primary mouse myoblast lines

Next, to look at the effects of VCP disease mutations instead of inhibition, primary myoblast lines were established from VCP^WT/WT^, VCP^R155H/WT^ and VCP^R155H/R155H^ mice [18]. Myoblasts were expanded and differentiated into mature myotubes for five days then treated with 1 mM sodium arsenite for 1 hour. The arsenite was removed and cells were washed with 1x PBS and replaced with fresh differentiation medium. Cells were harvested at specific recovery timepoints. While there appeared to be a slight increase in the amount of insoluble TDP-43 formed immediately following arsenite treatment (0h of recovery), there also was a decreased clearance of insoluble TDP-43 at 4 hours of recovery in the heterozygous and homozygous lines. This supports a loss of function in VCP muscle cells, again in endogenous levels of both TDP-43 and VCP.

### VCP inhibition slows aggregate clearance in seeded TDP-43 FRET assay

Next, to look at VCP inhibition in aggregates that were formed through proteopathic seeding rather than inducible overexpression or arsenite stress, we next utilized the TDP-43 FRET biosensor line. These are HEK293 cells stably expressing c-terminal fragments of TDP-43 fused to mruby3 and mclover3. Upon aggregation, there is a shift in fluorescence that can be measured by flow cytometry. To induce aggregation in these cells, they were seeded with lysates from HSA-hTDP-43^ΔNLS^ mouse muscle. To specifically look at VCP involvement in aggregate clearance rather than the initial seeding process, the cells were treated with VCP inhibitor on day 5 post seeding. In this case, we used the inhibitor NMS-873 as we found that CB-5083 was too toxic on the cells. Following 24 hour treatment with NMS-873, there was increased FRET signal compared to control DMSO treated cells and even increased from the pre-treatment timepoint. This increase could be due to cell proliferation during the treatment period or further aggregate spread occurring due to the decrease in disaggregase activity caused by NMS-873.

### HSA-hTDP-43^ΔNLS^ x VCP^R155H/WT^ mice have decreased clearance of insoluble TDP-43

To investigate how VCP patient mutations may influence TDP-43 muscle pathology *in vivo*, we utilized our previously characterized HSA-hTDP-43^ΔNLS^ mice [17] and crossed them with heterozygous VCP^R155H/WT^ knock in mice [18]. Expression of the cytoplasmic mislocalized TDP-43 was induced with doxycycline chow for 2 weeks followed by a return to regular chow for aggregate clearance timepoints. Hindlimb muscles were harvested from HSA-hTDP-43^ΔNLS^ mice and HSA-hTDP-43^ΔNLS^ x VCP^R155H/WT^ mice at specific accumulation and clearance timepoints (**Fig. 5A**) and analyzed for TDP-43 pathology. There did not appear to be any differences between the two genotypes in the level of accumulation of TDP-43 aggregates after two weeks on dox chow as shown by immunostaining for pTDP-43 and western blotting for insoluble TDP-43/pTDP-43 (**Fig. 5B**,**C**). However, there was a persistence of aggregates in the HSA-hTDP-43^ΔNLS^ x VCP^R155H/WT^ mice compared to HSA-hTDP-43^ΔNLS^ only mice after 3 weeks of recovery. By immunostaining the HSA-hTDP-43^ΔNLS^ mice have no remaining pTDP-43 positive aggregates remaining but HSA-hTDP-43^ΔNLS^ x VCP^R155H/WT^ mice have scattered affected fibers (Fig. 5B). By western blot, there is a slight increase in insoluble TDP-43 seen in the HSA-hTDP-43^ΔNLS^ x VCP^R155H/WT^ mice compared to HSA-hTDP-43^ΔNLS^ mice at the 3 week recovery timepoint (**Fig 5C**, red box). This *in vivo* model of VCP mutation in the presence of muscle-specific TDP-43 aggregation indicates reduced disaggregase activity in the presence of VCP mutations.

**Figure 5.**
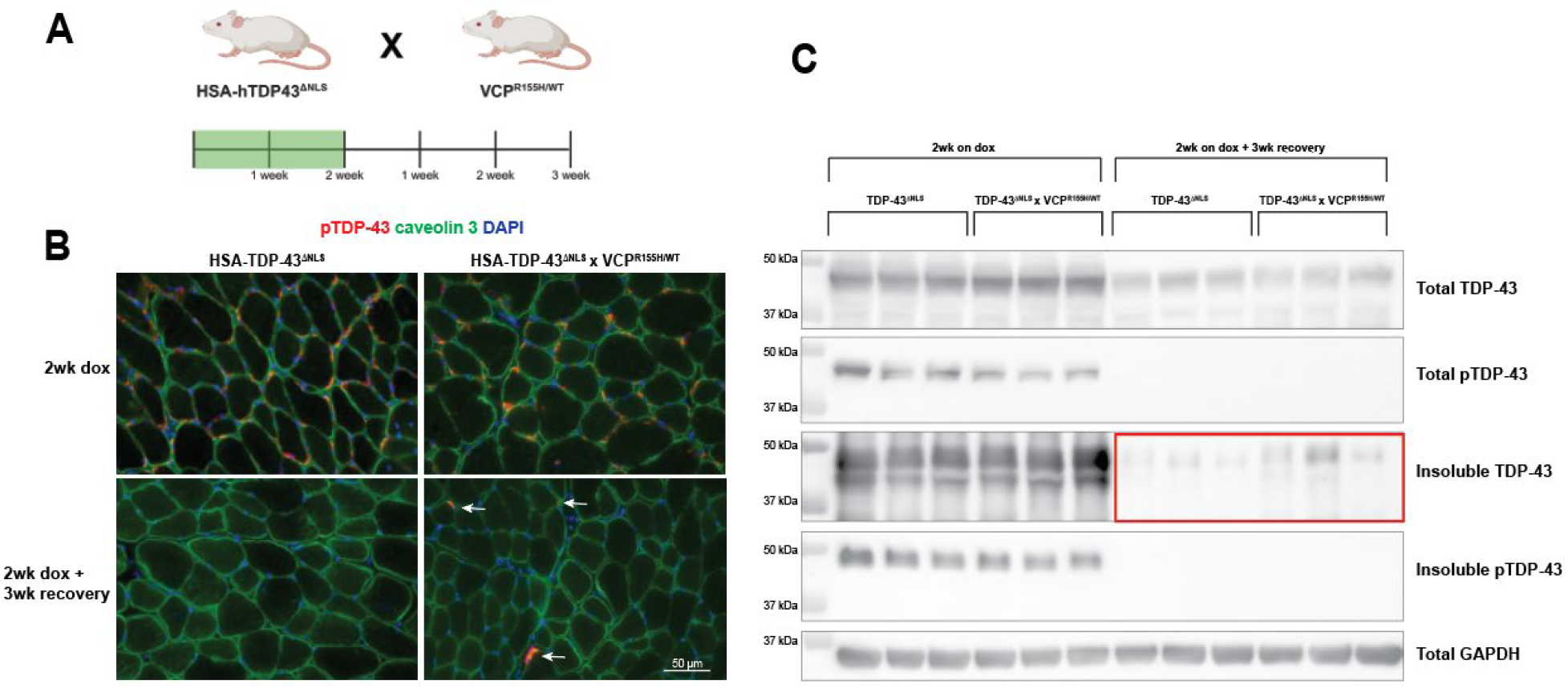
VCP mutant mice crossed with inducible muscle-specific TDP-43 aggregation mice have delayed clearance of TDP-43 aggregates. **(A)** HSA-hTDP-43^ΔNLS^ mice were crossed with VCP^R155H/WT^ heterozygous knock in mice, then treated with doxycycline chow for 2 weeks to activate the transgene and thus cause accumulation of sarcoplasmic aggregates. **(B)** While both genotypes show aggregate formation to a similar degree at two weeks on dox, at the three week recovery timepoint the HSA-hTDP-43^ΔNLS^ mice have minimal noticeable aggregates while there are still some sparse aggregates remaining in the HSA-hTDP-43^ΔNLS^ x VCP^R155H/WT^ mice. **(C)** Western blotting of the RIPA insoluble fraction from both genotypes again shows similar levels of accumulation of insoluble TDP-43 but a delayed clearance in the mice crossed with VCP mutant mice at three weeks of recovery (red box).

## Discussion

We demonstrated that VCP is critical for the clearance of insoluble TDP-43 in skeletal muscle using a variety of *in vitro* and *in vivo* model systems. In these systems, the R155H mutation in VCP appears to cause a loss of function similar to VCP inhibition with CB-5083 or NMS-873. However, this is only one of many patient mutations which may have different effects on TDP-43 pathological changes. There is evidence in support of both gain of function and loss of function effects of VCP patient mutations. Increased VCP ATPase activity has been noted from several mutations including R155H [20-24]. Additionally, VCP inhibitors or antisense oligonucleotide reduction of VCP were beneficial in animal models of VCP-related pathology [25-27]. Therefore, it has been proposed to inhibit or reduce VCP as a therapeutic option. In support of a loss of function in VCP disease, studies in which VCP was conditionally inactivated in the skeletal muscle or brain of adult mice recapitulated aspects of VCP pathology seen in IBM and FTD respectively [28, 29]. More recently, VCP activating drugs have been developed which aim to enhance the clearance of aggregated proteins [30, 31]. In particular, compounds that increase the D2 ATPase activity of VCP have shown beneficial effects in clearing insoluble TDP-43 in a cellular model of proteostatic stress featuring intranuclear TDP-43 aggregates [32].

Instead of thinking about VCP mutations as purely gain or loss of function, it is likely a much more nuanced situation, involving changes in cofactor interactions and downstream effects. We hypothesize that although ATPase activity is increased with many VCP mutations, at the same time there could be disrupted interactions with specific VCP cofactors causing a loss of disaggregase activity. Importantly, VCP has more than 30 different cofactor/adaptor proteins that it interacts with which help determine its cellular function [2]. VCP inhibitors may have different levels of potency when acting on VCP alone or VCP in a complex with specific other cofactors [33]. Perhaps rather than drugs that inhibit or enhance VCP ATPase activity, the next generation of drugs should be modulators of VCP-cofactor interactions. Therefore, our future work will look into the VCP cofactors involved and the mechanisms behind the dissolution of insoluble TDP-43 in different conditions (such as under cell stress versus seeded aggregate models). For example, one study found that soluble TDP-43 is primarily degraded by the proteasome but insoluble TDP-43 requires autophagy [34]. Additionally, a more recent paper found that VCP in combination with cofactors UFD1-NPL4 and FAF1 is involved in a “piecemeal autophagy” method of clearing aggresomes including those containing TDP-43 [35]. Future work will also utilize VCP patient iPSC-derived skeletal myocytes and neurons to study VCP cofactor involvement in TDP-43 disaggregase activity as perhaps some of the conflicting data so far has derived from the use of animal models which may have confounding differences from human VCP biology.

While we demonstrated in several model systems the importance of VCP in the resolution of insoluble TDP-43, there is still much to learn about the specific mechanisms behind VCP processing of TDP-43 and how to target these interactions for therapeutic interventions that may help reverse the disease course for patients with TDP-43 proteinopathies affecting both muscle and brain.

